# Computational Insights on Xenobiotic Nuclear Receptor-Air Pollutant Interactions Through Molecular Docking and Dynamics Simulations

**DOI:** 10.1101/2023.09.15.558018

**Authors:** Samee Ullah, Nasir Ali Khan

## Abstract

Air pollution is a major global health risk, contributing to millions of deaths each year. Exposure to air pollutants can lead to a variety of adverse health effects, including oxidative stress, inflammation, and DNA damage. The nuclear receptor pregnane X receptor (PXR) is a xenobiotic sensor that plays a key role in the detoxification of environmental chemicals and air pollutants can activate these receptors, leading to the expression of genes involved in drug metabolism and detoxification. Elucidating the structural interactions of air pollutants with PXR provides insights into the mechanisms by which these pollutants exert their toxic effects. Using computational techniques, we screened 124 air pollutants toxic compounds from (APDB) against PXR and identified priority compounds for molecular dynamics simulation that revealed the dynamic binding mechanisms and structural conformations involved in PXR activation. Furthermore, the root mean square deviation (RMSD) and root mean square fluctuation (RMSF) analyses characterized the stability and flexibility of PXR-pollutant complexes. Together, these computational approaches provide unprecedented insights into the molecular basis of PXR activation by air pollutants. By enabling high-throughput screening and multi-faceted biophysical characterization of PXR interactions, this work elucidates the structural mechanisms linking air pollution to impaired xenobiotic metabolism. Integration of virtual screening, molecular dynamics, and rigorous biophysical analyses represents a powerful computational systems pharmacology pipeline for assessing health risks posed by ubiquitous environmental toxicants.

## Introduction

Air pollution is one of the major environmental risks to health and is estimated to cause approximately 4.2 million premature deaths worldwide per year. Ambient (outdoor) air pollution alone contributes to around 3 million deaths annually [1]. The air pollutants implicated include particulate matter (PM), ozone (O3), nitrogen dioxide (NO2) and sulfur dioxide (SO2) which have both short-term and long-term health effects. Short-term exposure can lead to exacerbation of asthma, respiratory infections and increased hospital admissions [2]. Long-term exposure contributes to increased risk of cardiovascular morbidity such as stroke, heart disease, lung cancer, chronic obstructive pulmonary disease (COPD) as well as premature mortality [3].

Air pollutants elicit adverse health effects by inducing oxidative stress, inflammation, DNA damage and epigenetic alterations [4]. The nuclear receptors comprise a superfamily of ligand-activated transcription factors that regulate gene expression and are key players in maintaining cellular homeostasis [5]. The pregnane X receptor (PXR), also known as NR1I2, SXR or PAR, is a nuclear receptor that regulates genes involved in xenobiotic metabolism and acts as a xenobiotic sensor [6]. Upon activation by a ligand, PXR transcriptionally regulates broad specificity drug metabolizing enzymes like CYP3A4 and CYP2B6 and transporters like MDR1 involved in clearance and detoxification [7]. Air pollutants such as polycyclic aromatic hydrocarbons (PAHs) and dioxins have been identified as PXR activators and ligands [8]. The promiscuous nature of ligand binding makes PXR a molecular target for diverse environmental chemicals. Elucidating the structural mechanisms and dynamics of xenobiotic interactions with PXR can provide toxicological insights into air pollutant health impacts.

Computational methods like molecular docking and molecular dynamics (MD) simulations have emerged as powerful techniques complementary to experimental studies, providing atomistic insights into ligand-protein interactions [9]. Molecular docking predicts the bioactive conformation of small molecule ligands within the binding sites of target proteins. It allows rapid virtual screening of large chemical libraries [10]. MD simulations model the motions of atoms and molecules over time by integrating Newton’s equations of motion and enable investigating thermodynamics and conformational dynamics of protein-ligand complexes [11].

In this study, molecular docking of 124 air pollutants from the Air Pollutants Database (APDB) was performed with the pregnane receptor to determine their binding affinities and modes. This was followed by molecular dynamics simulations of selected top docked complexes from GlideXP(Extra-precision) to gain structural conformational dynamic perspectives into PXR-pollutant interactions. The findings from this study provide the binding mode and structural basis of air pollutant interactions with pregnane receptors (PXR). Using computational modeling and simulations, we have determined the structure of the xenobiotic receptor PXR ligand-binding mode and characterized the molecular determinants of pollutant binding. Our simulations reveal the binding mode for various air pollutants toxins and provide structural insights into the molecular mechanisms of PXR activation. By enabling prediction of PXR-pollutant interactions, these computational biophysical techniques allow identification of health-impacting pollutants for prioritized toxicological assessment. Modeling of the PXR ligand-binding pocket and dynamics establishes a foundation for screening large libraries of existing and emerging pollutants for interactions with PXR signaling involved in xenobiotic clearance. Our computational approach sheds light on the complex connections between air pollution and impaired xenobiotic metabolism mediated by nuclear receptors. Integration of computational simulations with an understanding of receptor signaling mechanisms holds promise for elucidating the health risks posed by ubiquitous environmental pollutants.

## Materials and Methods

The complete workflow of the current study is provided in Figure 1.

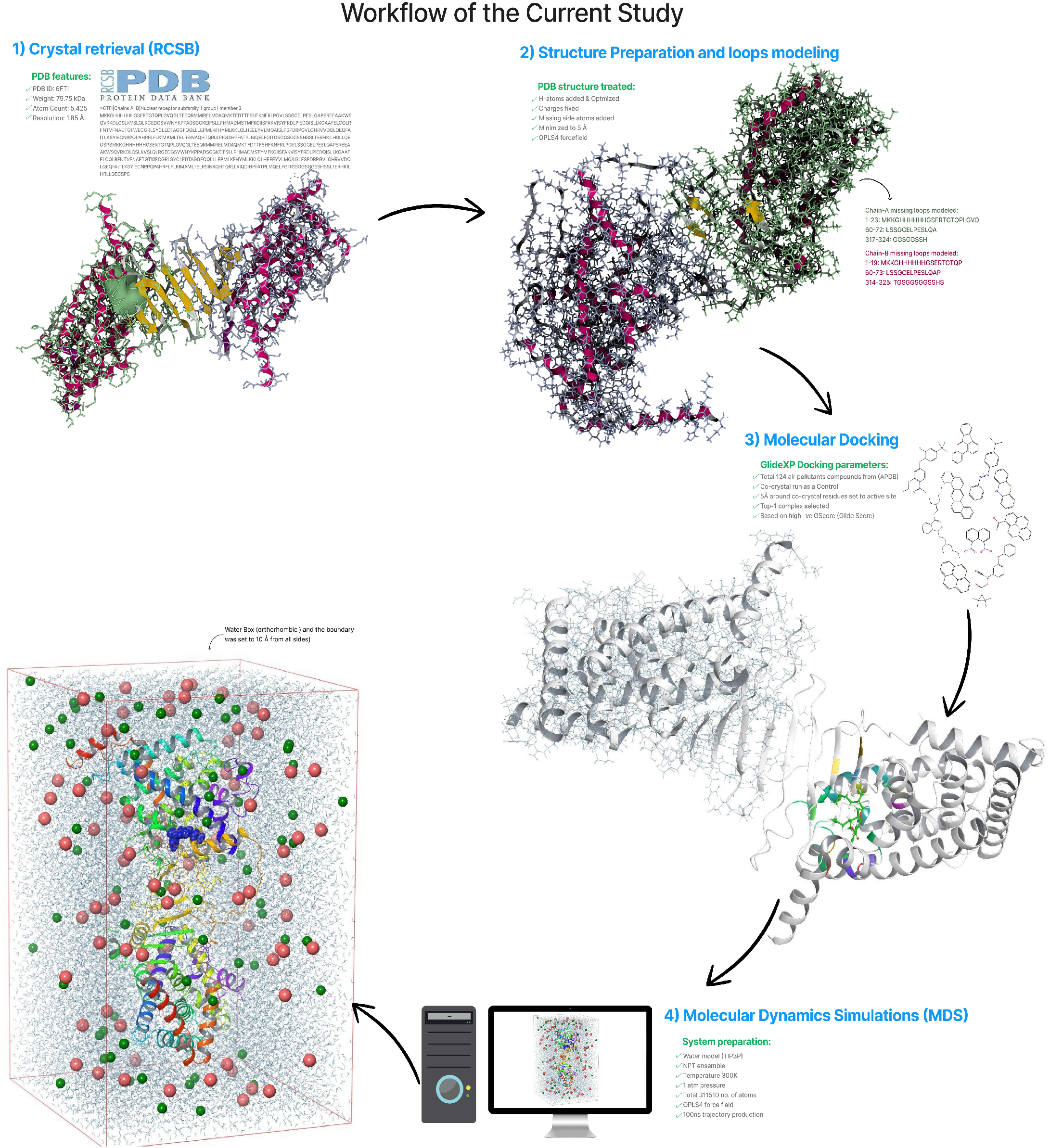

### Crystal Structure Retrieval and Preparation

The crystal structures of the PXR protein (PDB ID: 6TFI) and the air pollutants compounds of interest herein total 124 were retrieved from the Protein Data Bank (PDB) and Database of Air Pollutants (APDB) [12,13]. The PDB is a database of experimentally determined protein structures while the APDB is the database on air pollutants characterization and similarity prediction provided by the Environmental Protection Agency (EPA). The waters and ions were removed from crystal structure. The hydrogen and side chain atoms were added and optimized. The bond orders corrected, and further energy minimized to 5 Å using the OPLS4 force field [14]. The neutral terminal caps n-methyl amide (NME) and acetyl (ACE) were added at both N and C terminal of the structures. This structure was further used for molecular docking and molecular dynamics simulations. The ligands minimization is achieved by converting the canonical smiles in to 3d structures using the LigPrep and the artifacts in the structures checked and removed.

### Missing Loops Modeling

In chain A of the protein structure, residues 1-23 (MKKGHHHHHHGSERTGTQPLGVQ), 60-72 (LSSGCELPESLQA), 317-324 (GGSGGSSH), and the C-terminal proline-serine (PS) were missing in the native crystal structure shown in Figure 2. In chain B, residues 1-19 (MKKGHHHHHHGSERTGTQP), 60-73 (LSSGCELPESLQAP), 314-325 (TGSGGSGGSSHS), and the C-terminal serine at position 344 were also missing. These missing regions were modeled using the loop modeling algorithm in molecular operating environment with the amber14 force field. The missing loops were modeled based on homology search using the loop modeling engine of molecular operating environment. The loops were searched using the PDB with RMS limit to 0.5 Å, loop limit 1000 and energy window 10 k/cal mol. In all cases the best loops were selected based on highest coarse score (Egeom: Phi, Psi Ramachandran energy, EvdW: Vander Waals interaction loop energy and Ehb: Hydrogen bond interaction loop energy). The modeled loop’s structure was superposed on to the crystal checked and analyzed in figure 3.

**Figure 2.**
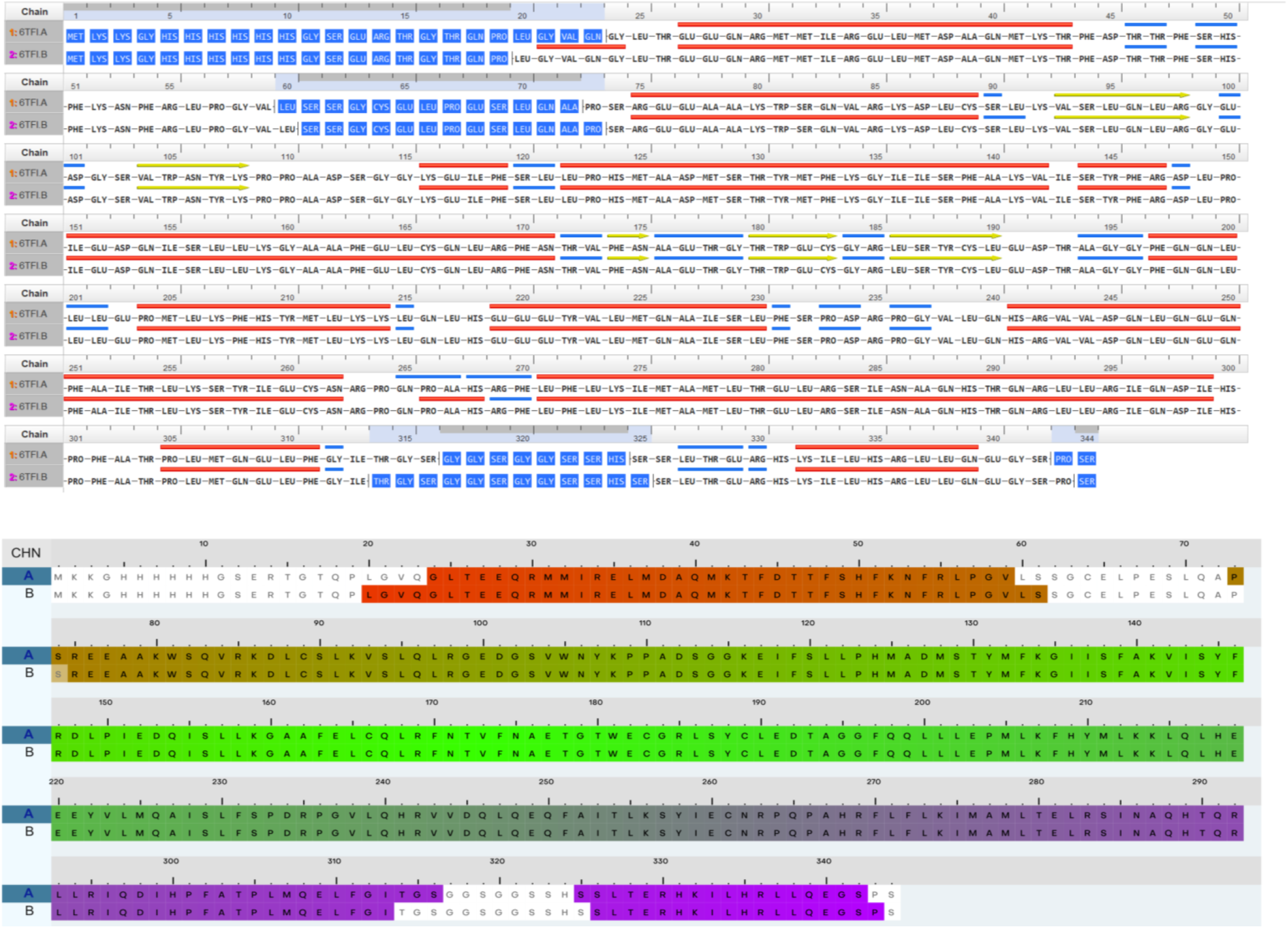
The unresolved residues in the crystal structure highlighted in blue color in both chains.

**Figure 3.**
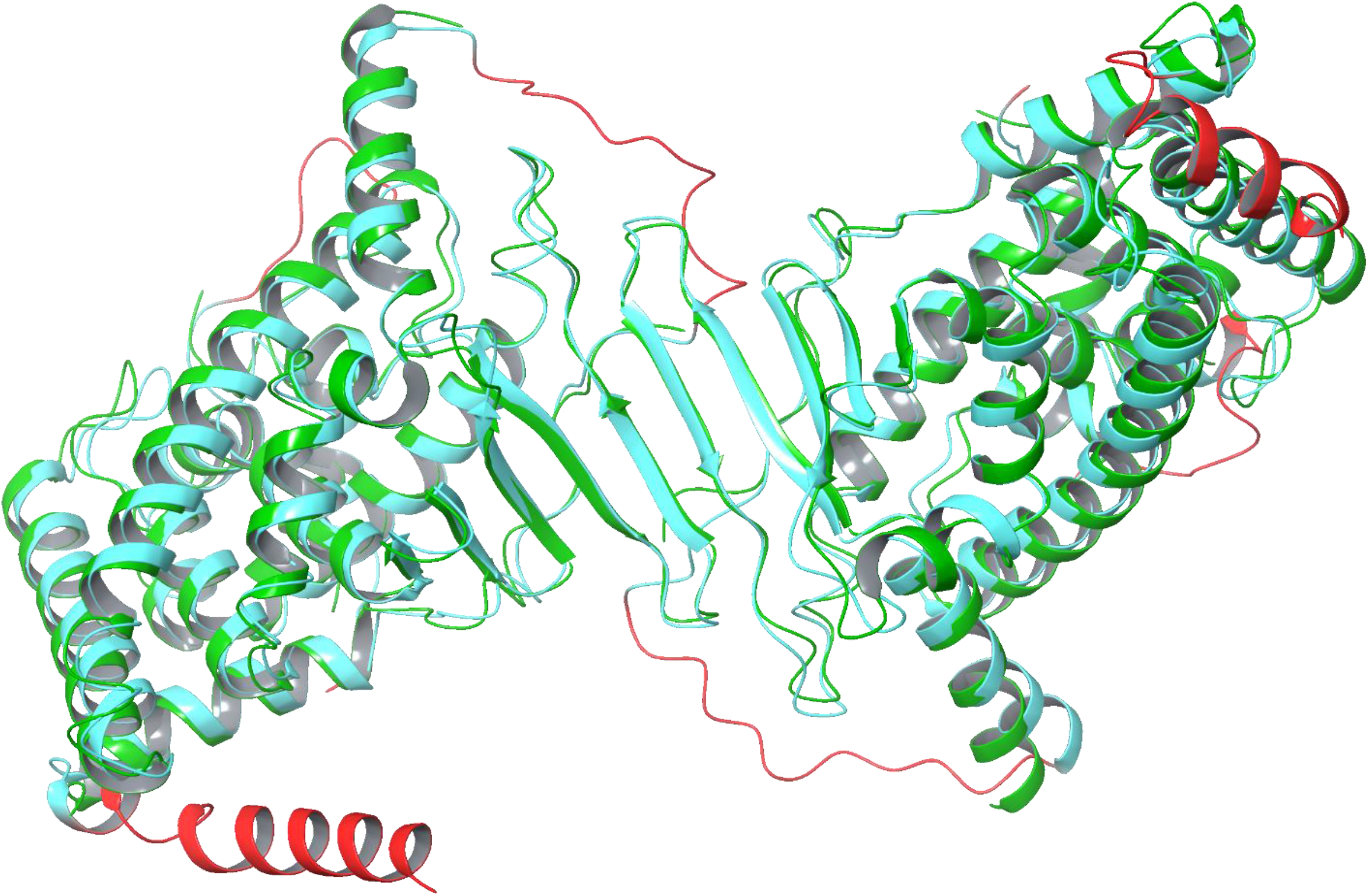
The modeled loops superposed and colored in red. The green color highlighted is the native structure and the cyan after loops modeled.

### Receptor Grid Generation and Docking

A docking grid was generated around the active site of the co-crystal ligand in chain b structure to represent the space where the air pollutants small molecules could bind. The default grid Van der Waals radius scaling is set to 1.0 scaling factor and 0.25 partial charge cutoff. The enclosed box is set to 20 Å at all sides and the ligand diameter midpoint box in green to 10 Å in figure 4. The grid coordinates of X=51.53 Y=36.61 and Z=15.02 were also recorded. The PXR (pregnane X receptor) and the 124 small molecules compounds imported into glide docking engine. The default scaling Vander Waals radii of 0.80 scaling factor and partial charge cutoff 0.15 and the precision is set to XP (Extra-Precision) [15]. The ligand sampling is set to flexible and sample nitrogen inversions, ring conformations and Epik state penalties to docking score were enabled [16]. Finally, the docking was run on Fedora 38 Scientific OS (operating system) using the AMD Ryzen 9 CPU (central processing unit) with 64 GB (gigabyte) of RAM (random access memory). The top-scoring docked complexes were further analyzed.

**Figure 4.**
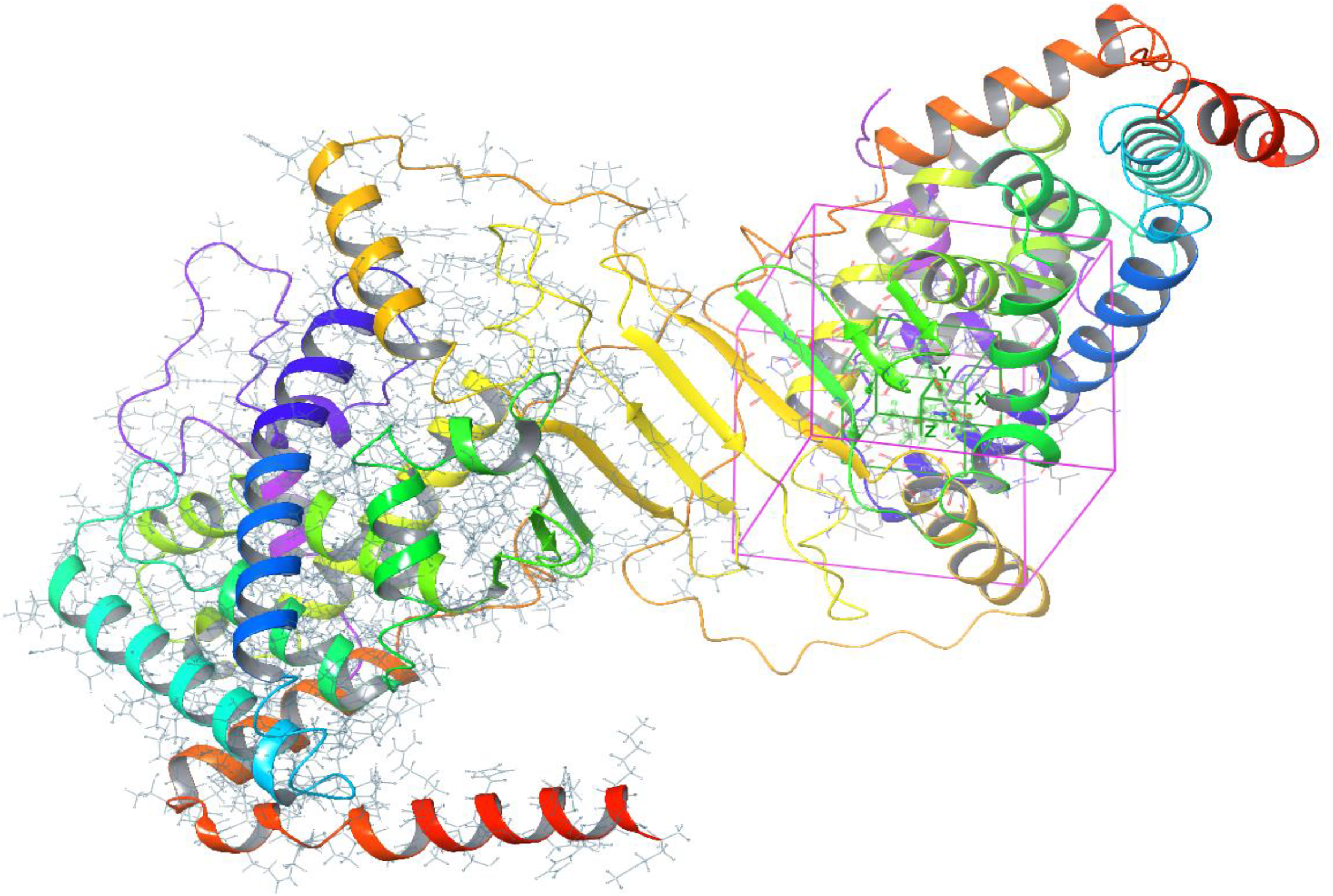
Receptor grid generated based on the co-crystal ligand coordinates at the chain b structure.

### Control Docking

In this study, the co-crystallized ligand (ligand ID: N6H) was also docked into the receptor grid to serve as a control. N6H is a small molecule with formula (C_22_H_24_C_l_N_3_O_4_) and molecular weight of (429.897 g/mol) that binds to the PXR protein. The binding affinity of the co-crystallized ligand was used to assess the accuracy of the docking results. Binding affinity is a measure of the strength of the binding interaction between a ligand and a receptor. The higher the binding affinity, the stronger the binding interaction.

### Molecular Dynamics Simulations (MDS)

The top-scoring docked complexes were subjected to molecular dynamics (MD) simulations using the Desmond[17]. MD simulations allow the study of the dynamic interactions between the small molecule ligands and the protein structures. The MD simulations were carried out for a period of 50 nanoseconds. The top1 complex imported into the orthorhombic box and solvated with TIP3P model [18]. The system neutralized by adding 02 Na ions and the salt concentration to 0.1M NaCl. The system’s atoms 93211 and volume 1819442 checked and recorded. The simulation time is set to 50ns (nano second) and recording interval 50ps (pico-second). The approximate number of frames is set to 1000 trajectory frames. The temperature is set to 300K(kelvin) and the pressure 1.0 (bar). Finally, the system relaxation achieved by the default relax model before simulation protocol and the 50ns trajectory run performed using the OPLS4 force field.

## Results

### Docking Interaction Analysis

To identify potential ligands for pregnane X receptor (PXR), we performed docking analysis of 124 compounds from the ABDB database using the glide-XP algorithm. We selected the top 5 complexes based on the highest glide binding energy, which is a measure of the strength of the interaction between the ligand and the receptor. The more negative the glide binding energy, the stronger the binding affinity. Table-1 shows the top 5 complexes and their corresponding glide binding energy values. The top-ranked compound (Molecule_000008) had a glide binding energy of -43.95 kcal/mol, indicating that it was the most potent binder for PXR. The other four compounds (Molecule_000085, Molecule_000032, Molecule_000043, Molecule_000124) had glide binding energies of -39.16, -39.43, -28.30, and -35.38 kcal/mol, respectively, suggesting that they were also good binders for PXR.

**Table 1.**
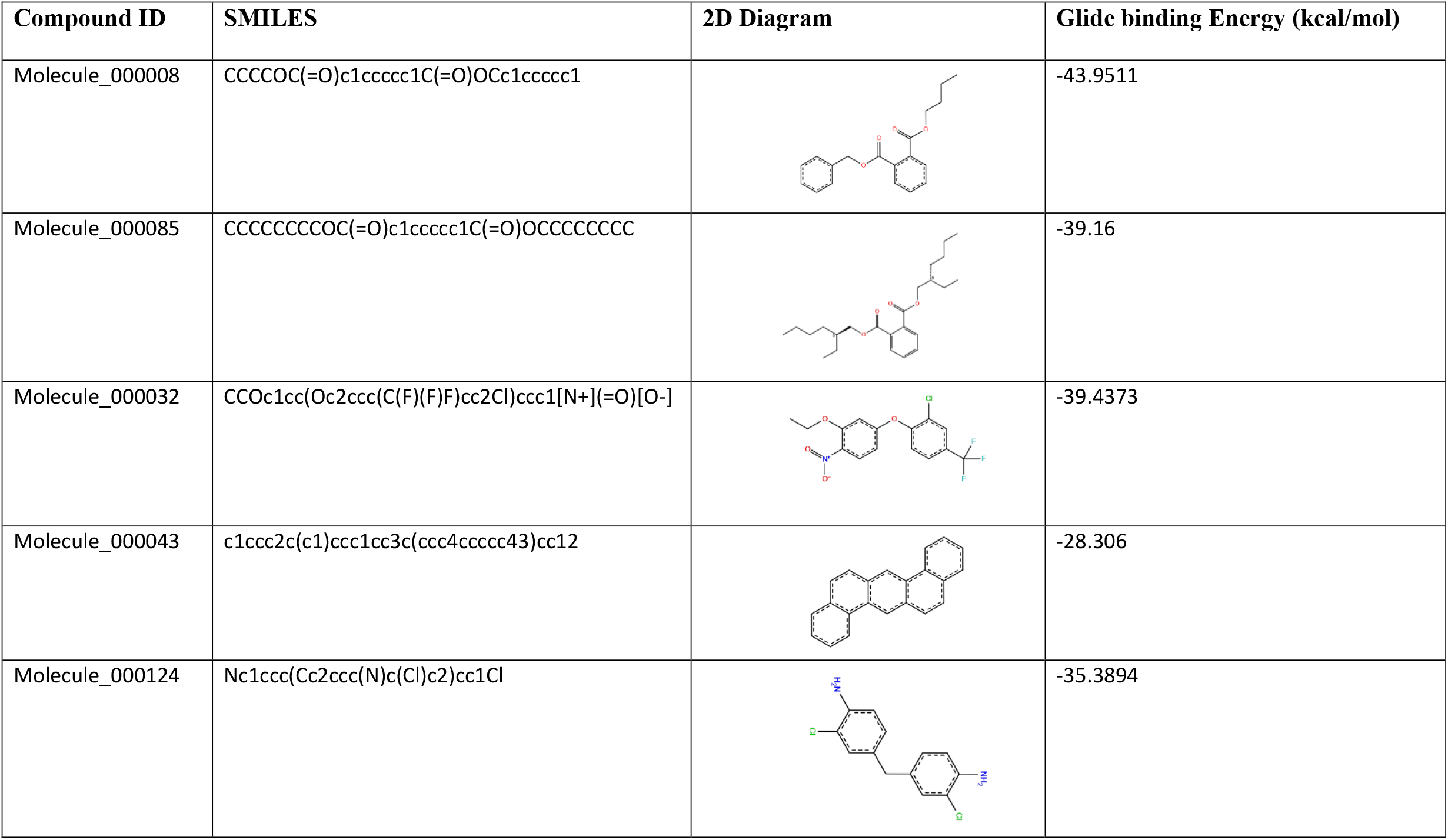
illustrates the glide binding energy of top-05 complexes predicted via Glide-XP algorithm.

| Compound ID | SMILES | 2D Diagram | Glide binding Energy (kcal/mol) |
| --- | --- | --- | --- |
| Molecule_000008 | CCCCOC(=O)c1ccccc1C(=O)OCc1ccccc1 |  | -43.9511 |
| Molecule_000085 | CCCCCCCCOC(=O)c1ccccc1C(=O)OCCCCCCCC |  | -39.16 |
| Molecule_000032 | CCOc1cc(Oc2ccc(C(F)(F)F)cc2Cl)ccc1[N+](=O)[O-] |  | -39.4373 |
| Molecule_000043 | c1ccc2c(c1)ccc1cc3c(ccc4ccccc43)cc12 |  | -28.306 |
| Molecule_000124 | Nc1ccc(Cc2ccc(N)c(Cl)c2)cc1Cl |  | -35.3894 |

To further investigate the binding mode and interactions of these compounds with PXR, we analyzed their 2D and 3D structures binding mode interactions. We found that all five compounds formed hydrogen bonds and/or pi-pi stacking interactions with key residues in the active site of PXR. These interactions are important for stabilizing the ligand-receptor complex and enhancing the specificity of binding. Figure 5 shows the 2D and 3D structures of Molecule_000008 bound to PXR. This compound formed a single hydrogen bond of 2.14 Å distance between its O7-H9529 atom and the side chain nitrogen atom of GLN167 residue in PXR. This residue is conserved among different species and plays a crucial role in ligand recognition and activation of PXR. Figure 6 shows the 2D and 3D structures of Molecule_000085 bound to PXR. This compound formed a single hydrogen bond of 2.0 Å distance between its O10-H9529 atom and the side chain nitrogen atom of GLN167 residue in PXR. In addition, it also formed a pi-pi stacking interaction with HIS289 residue in PXR. This residue is located at the bottom of the active site and contributes to the hydrophobic environment that favors ligand binding. Figure 7 shows the 2D and 3D structures of Molecule_000032 bound to PXR. This compound formed a single hydrogen bond of 2.08 Å distance between its O19-H9529 atom and the side chain nitrogen atom of GLN167 residue in PXR. Figure 8 shows the 2D and 3D structures of Molecule_000043 bound to PXR. This compound formed two pi-pi stacking interactions with TRP181 and HIS289 residues in PXR.

**Figure 5.**
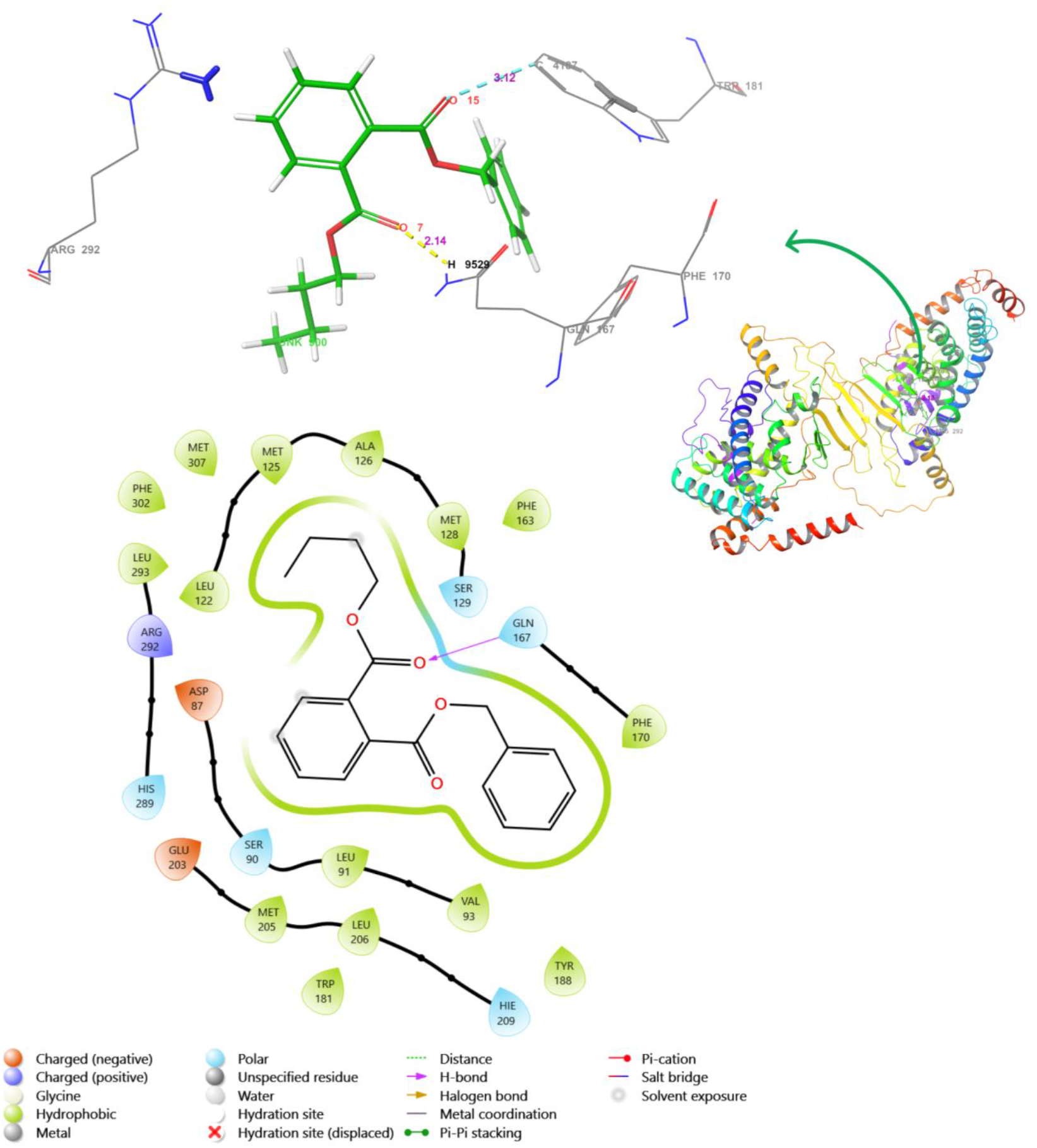
illustrates the 3D and 2D interaction diagram of top1-complex. The top1 ligand Molecule_000008 is depicted in green and the interacting residues in grey. The H-bond in yellow dash line and distance in salmon pink.

**Figure 6.**
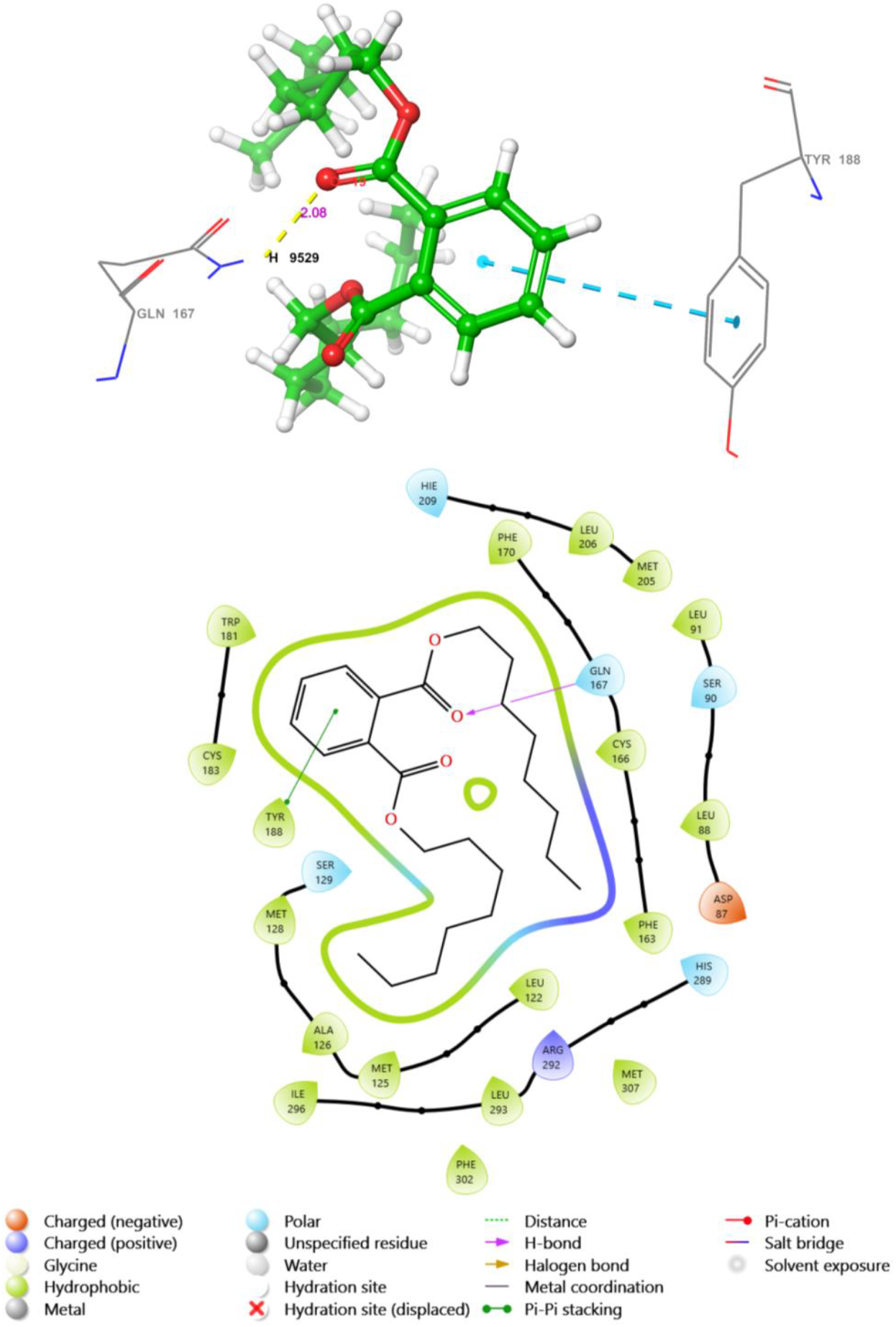
illustrates the 3D and 2D interaction diagram of top2-complex. The ligand Molecule_0000085 is depicted in green and the interacting residues in grey. The H-bond in yellow dash line and distance in salmon pink.

**Figure 7.**
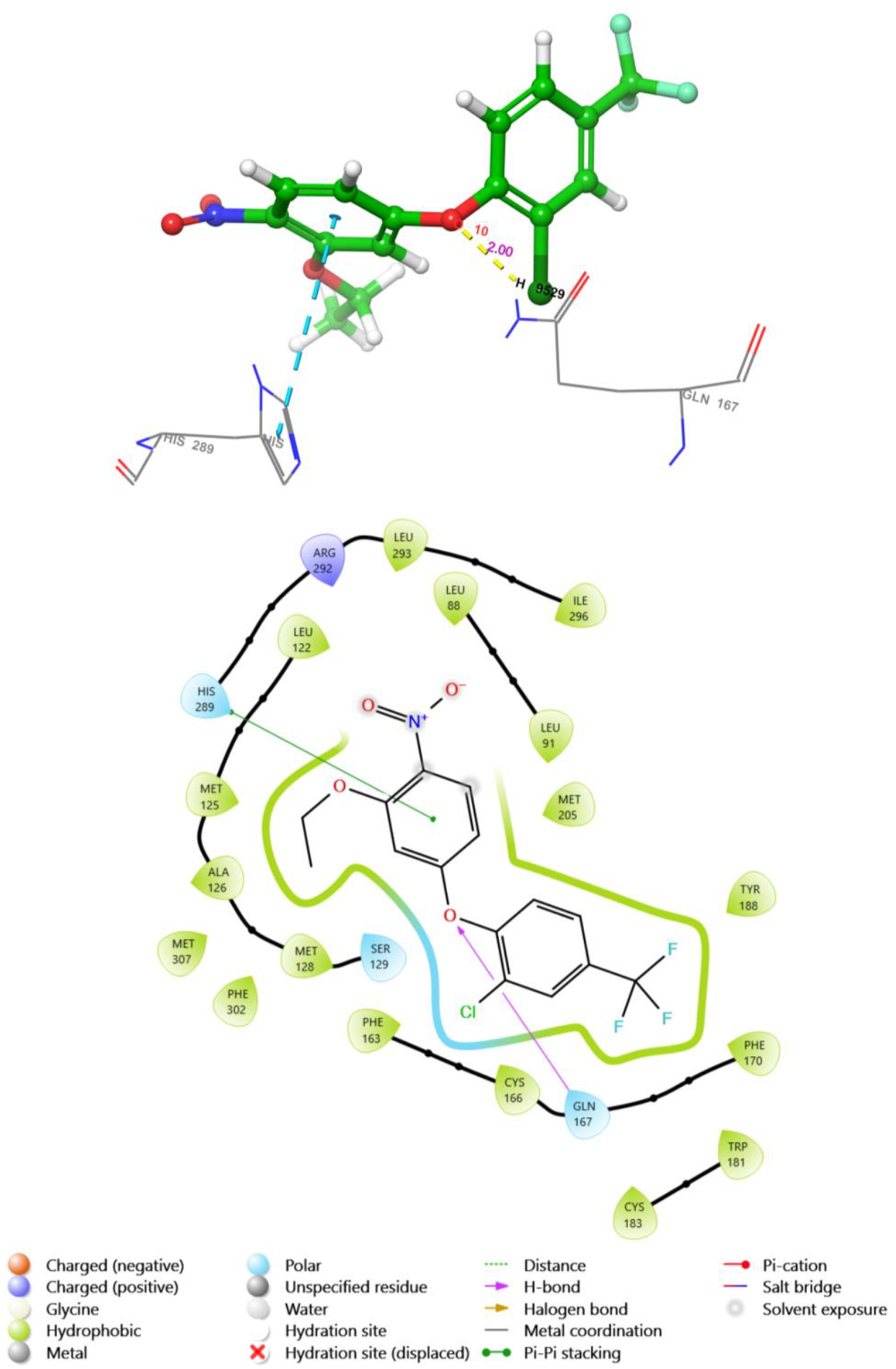
illustrates the 3D and 2D interaction diagram of top3-complex. The ligand Molecule_0000032 is depicted in green and the interacting residues in grey. The H-bond in yellow dash line and distance in salmon pink

**Figure 8.**
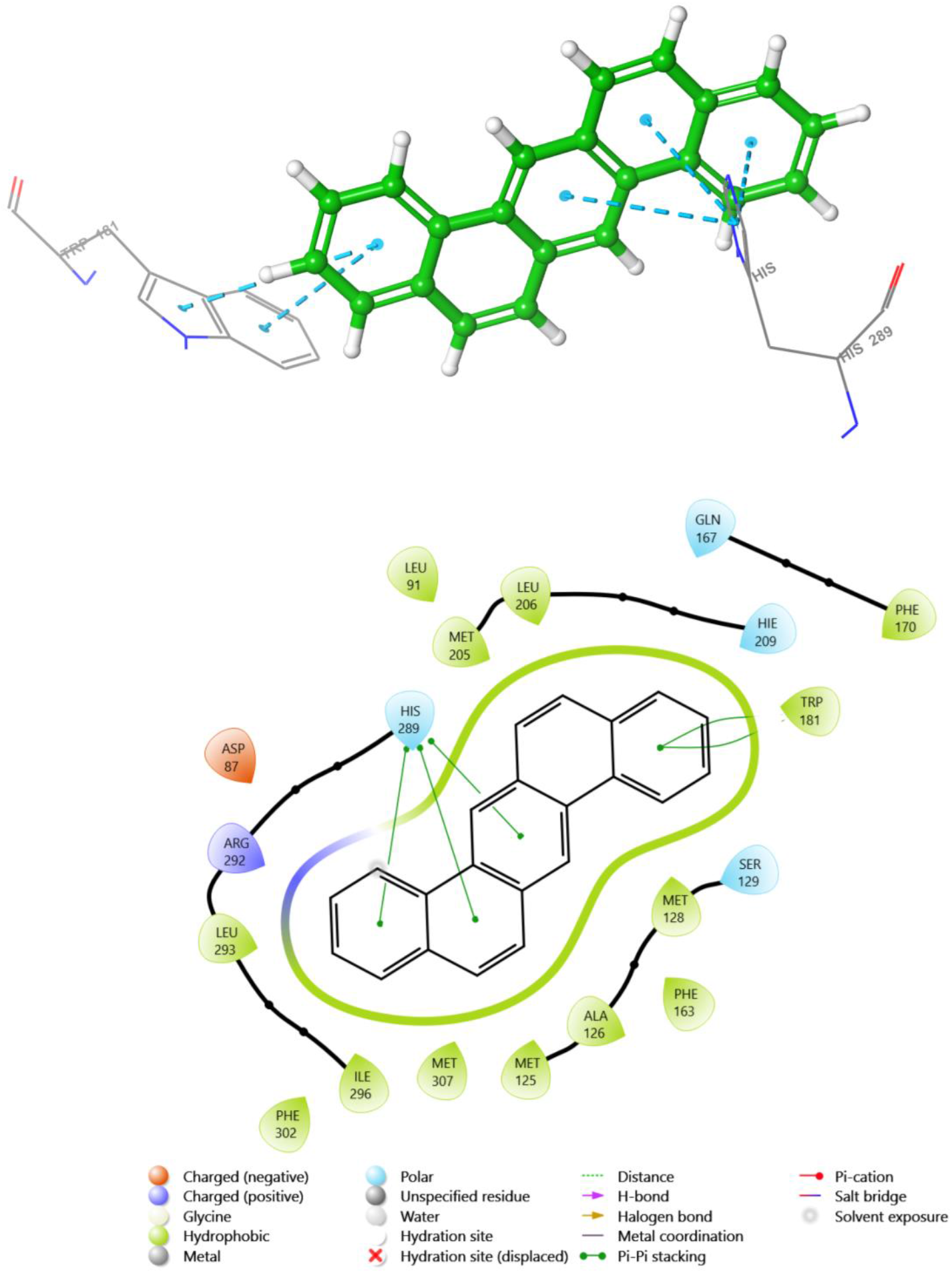
illustrates the 3D and 2D interaction diagram of top4-complex. The ligand Molecule_0000043 is depicted in green and the interacting residues in grey. The H-bond in yellow dash line and distance in salmon pink

Figure 9 shows the 2D and 3D structures of Molecule_000124 bound to PXR. This compound formed a pi-pi stacking interaction with PHE170 residue in PXR. PHE170 is also a conserved residue that forms part of the active site and interacts with different classes of ligands. Based on these results, we concluded that Molecule_000008 was the most promising candidate for further evaluation as a potential modulator of PXR activity. Therefore, we performed molecular dynamics simulation to assess its conformational stability and dynamics in complex with PXR over time.

**Figure 9.**
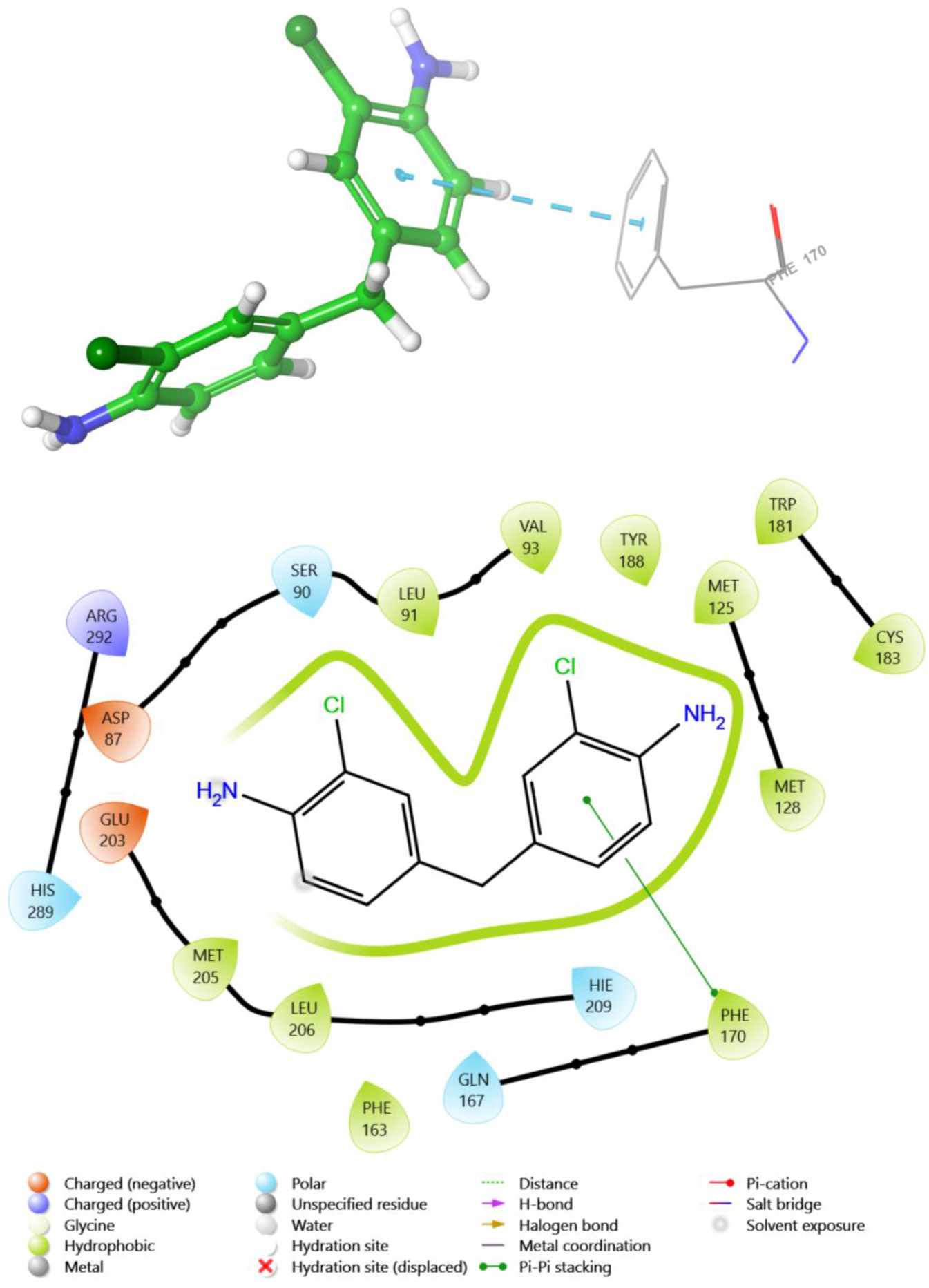
illustrates the 3D and 2D interaction diagram of top5-complex. The ligand Molecule_000124 is depicted in green and the interacting residues in grey. The H-bond in yellow dash line and distance in salmon pink

### PXR-top1 complex conformational landscape

The molecular dynamics (MD) simulations of the pregnane X receptor (PXR) complex were performed for 50 ns to comprehensively characterize the conformational dynamics, stability, and equilibrium behavior of the system. The protein backbone root mean square deviation (RMSD) relative to the crystal structure coordinates was calculated over the full trajectory to quantify the structural changes and flexibility over time (Fig. 10). During the initial 10 ns of the simulation, the RMSD showed a rapid increase from the crystal structure, reflecting structural relaxation as the complex evolved away from the static experimental model into a more native equilibrium state. By approximately 10 ns, the RMSD leveled off, forming a plateau that was maintained for the remainder of the 50 ns trajectory. This stable plateau region indicates an equilibrated ensemble representing the native state was achieved. The average RMSD value of the plateau was 4.30 ± 0.37 Å. However, close inspection of the RMSD plateau revealed local fluctuations periodically spiking above the baseline, reaching maximum values up to 5.43 Å before reverting back down. These temporary excursions reflect the intrinsic flexibility of PXR, with the receptor transiently sampling higher RMSD conformations before rapidly returning to the global minimum. Given the small amplitude of the spikes, these dynamics likely represent localized structural rearrangements rather than widespread unfolding events. Given the small amplitude of the spikes, these dynamics likely represent localized structural rearrangements rather than widespread unfolding events. The broad distribution of RMSD values followed a Gaussian shape centered at 4.30 Å. The single peak confirms PXR samples one major basin, without transitions between multiple distinct states over the 50 ns trajectory. Taken together, the rapid convergence, stable RMSD baseline, and unimodal distribution verify proper equilibrium behavior was attained.

**Figure 10.**
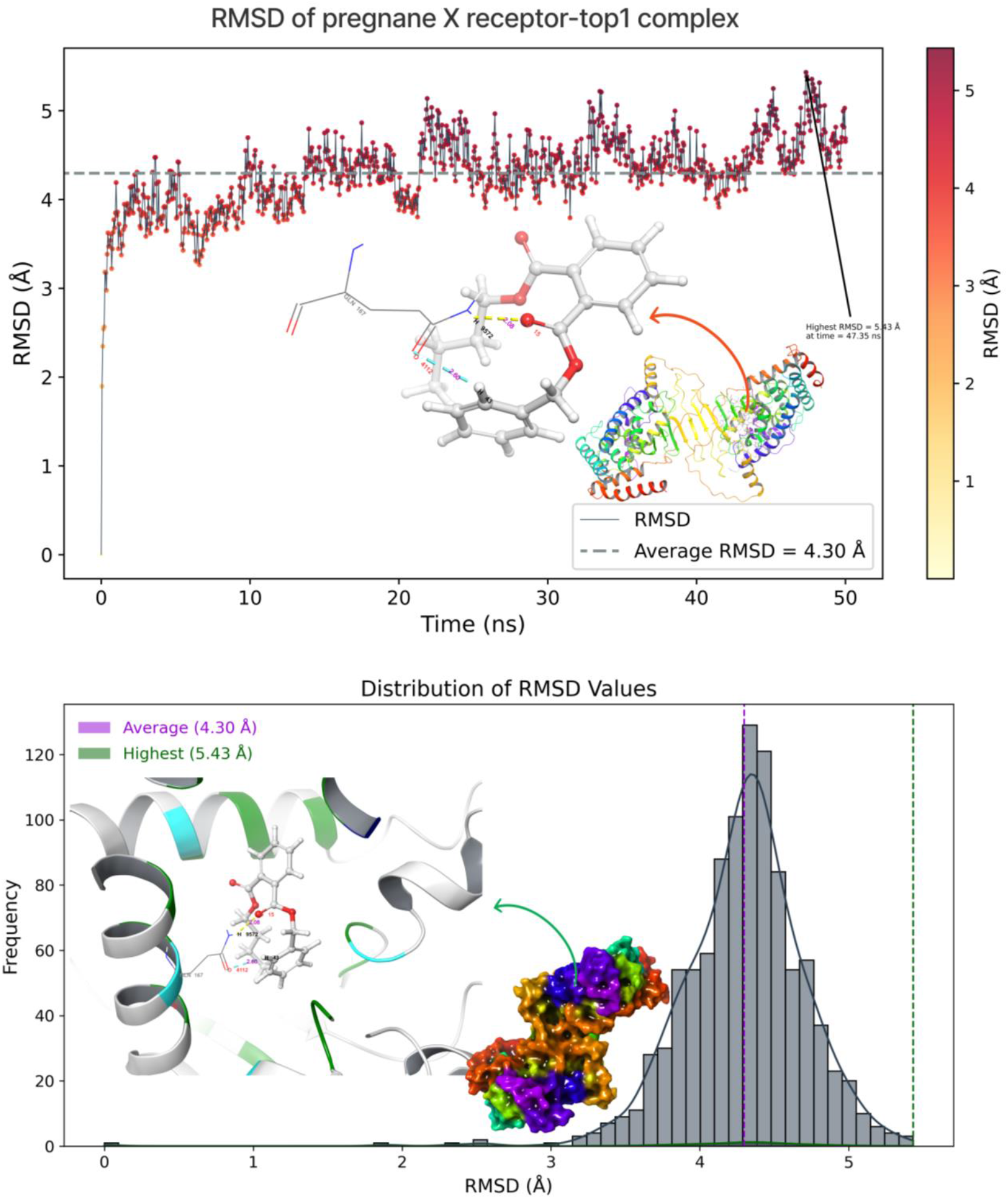
RMSD of pregnane X receptor top1-complex over 50 ns. The 5.4 Å peak (arrow) at 47.35 trajectory frame indicates a high rmsd (green dash line in histogram) and the 4.3 Å average (gray dashed line and pink in histogram).

Notably, the average RMSD of 4.30 Å is significantly higher than typical values for globular proteins, indicating PXR inherently explores a much more open, flexible conformational landscape. This likely facilitates its promiscuous nature, enabling accommodation of diverse ligands within the binding pocket. The large RMSD spikes highlight the ability to access expanded conformations while preserving the binding-competent ground state. The intrinsic mobility and structural plasticity are essential to biological function.

The RMSF plot in figure 11 shows the residue-wise root mean square fluctuation (RMSF) values for the protein complex over the trajectory. Higher RMSF values indicate more flexibility and mobility of those residues during the simulation. The N-terminal residues of chain A (residues 1-25) exhibit very high RMSF values, indicating this region is highly flexible and dynamic. This likely represents the disordered tail region that is not part of the structured domain. Similarly, the C-terminal residues of chain B (residues 340-344) also show high RMSF of 3.1 Å to 6.4 Å reflecting high mobility of this disordered C-terminal tail. Another major region of high flexibility is the loop region spanning residues 190-200. This loop shows RMSF values over 4Å, suggesting it samples a wide range of conformations during the simulation. Being a surface exposed loop, such mobility is expected. This plasticity likely facilitates binding interactions with ligands or other proteins. The catalytic cleft region spanning residues 316-319 also shows moderately high RMSF values between 5-6Å. Besides these major flexible regions, much of the protein core shows relatively low RMSF values under 2Å. This indicates these areas maintain a stable folded conformation during the simulation. The limited dynamics in the structured regions is expected for a properly folded native state. The RMSF analysis reveals protein segments with high internal mobility likely important for biological function, as well as more rigid structured regions that provide overall stability.

**Figure 11.**
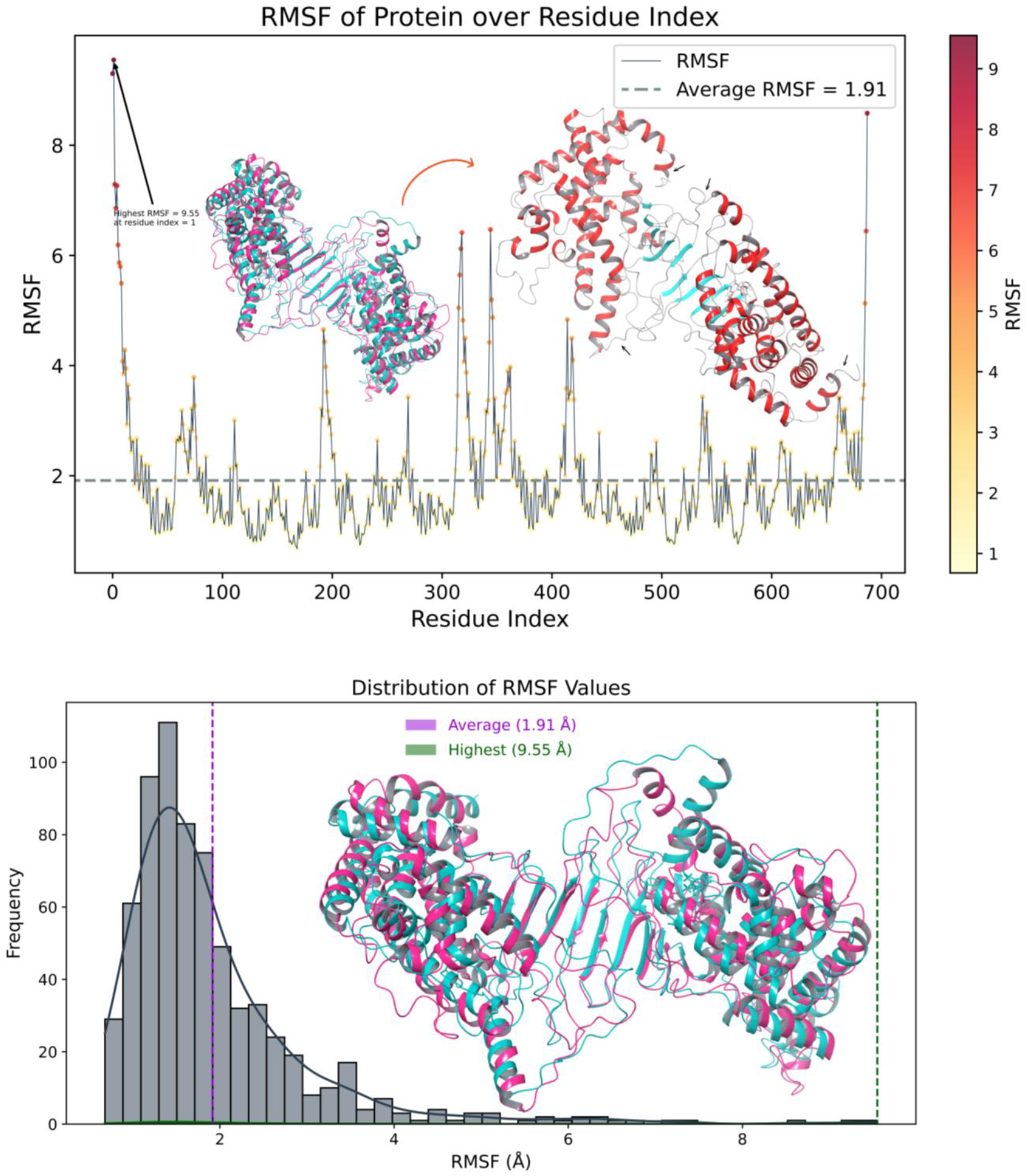
RMSF of pregnane X receptor residues in the top1-complex over 50 ns. The 9.5 Å peak (arrow) indicates a highly flexible C-terminus (green in histogram). Most residues exhibit RMSF near the 1.9 Å average (gray dashed line). A rigid 200-500 core has RMSF below 1 Å (pink in histogram).

## Conclusion

This study still in preliminary, provides valuable computational insights into the interactions between xenobiotic nuclear receptor pregnane X (PXR) and air pollutants obtained from APDB (Air Pollutant Database), shedding light on the binding mode, structural dynamics and mechanisms underlying the toxic effects of air pollution and sensing by nuclear receptor. Employing a comprehensive computational approach, including molecular docking and molecular dynamics simulations, we systematically screened 124 air pollutants to assess their interactions with PXR. The results revealed several compounds with high binding affinities to PXR, suggesting their potential as activators of this nuclear receptor. Furthermore, our molecular dynamics simulations probed the dynamic behavior of PXR-pollutant complexes over a 50-nanosecond period, shedding light on the inherent flexibility of the PXR protein, a crucial feature for its role in xenobiotic metabolism. The analysis showed that the PXR protein exhibits inherent flexibility and can access a broad range of conformations, which likely contributes to its ability to interact with diverse ligands. This structural plasticity is crucial for its role in xenobiotic metabolism. Furthermore, the study highlighted specific regions of the PXR protein that exhibited high flexibility, including N-terminal and C-terminal residues, as well as loop regions. These flexible regions are likely essential for binding interactions with pollutants, emphasizing their significance in PXR’s biological function. This research represents a significant contribution to the understanding of how air pollutants interact with PXR and impact xenobiotic metabolism. The computational approaches employed in this study provide a powerful tool for high-throughput screening of potential health-impacting pollutants and offer insights into the structural mechanisms connecting air pollution to impaired xenobiotic metabolism.

## Author contributions

SU planned the study and wrote the paper, modeled loops, performed and analyzed docking and molecular dynamics simulations (MDS), submitted the modeled structures to Protein Model Database (PMDB) and to Model Archive. NAK helped in the introduction of the manuscript.

## Data availability

The modelled loops of the structure in this study can be accessed at Protein Model Database PMDB ID: PM0084591 and Model Archive ID: ma-1iksv.

## References

[1] Ambient (outdoor) air pollution, (n.d.). https://www.who.int/news-room/fact-sheets/detail/ambient-(outdoor)-air-quality-and-health (accessed September 9, 2023).

[2] P. Zhang, G. Dong, B. Sun, L. Zhang, X. Chen, N. Ma, F. Yu, H. Guo, H. Huang, Y.L. Lee, N. Tang, J. Chen, Long-term exposure to ambient air pollution and mortality due to cardiovascular disease and cerebrovascular disease in Shenyang, China, PLoS One. 6 (2011) e20827. 10.1371/journal.pone.0020827.

[3] D.E. Newby, P.M. Mannucci, G.S. Tell, A.A. Baccarelli, R.D. Brook, K. Donaldson, F. Forastiere, M. Franchini, O.H. Franco, I. Graham, G. Hoek, B. Hoffmann, M.F. Hoylaerts, N. Künzli, N. Mills, J. Pekkanen, A. Peters, M.F. Piepoli, S. Rajagopalan, R.F. Storey, E.A. for C.P. and R. and E.H.F.A. on behalf of ESC Working Group on Thrombosis, Expert position paper on air pollution and cardiovascular disease, European Heart Journal. 36 (2015) 83–93. 10.1093/eurheartj/ehu458.

[4] D.M. DeMarini, Genotoxicity biomarkers associated with exposure to traffic and near-road atmospheres: a review, Mutagenesis. 28 (2013) 485–505. 10.1093/mutage/get042.

[5] R.M. Evans, D.J. Mangelsdorf, Nuclear Receptors, RXR, and the Big Bang, Cell. 157 (2014) 255–266. 10.1016/j.cell.2014.03.012.

[6] S.A. Kliewer, B. Goodwin, T.M. Willson, The nuclear pregnane X receptor: a key regulator of xenobiotic metabolism, Endocr Rev. 23 (2002) 687–702. 10.1210/er.2001-0038.

[7] C. Zhou, S. Verma, B. Blumberg, The steroid and xenobiotic receptor (SXR), beyond xenobiotic metabolism, Nucl Recept Signal. 7 (2009) e001. 10.1621/nrs.07001.

[8] J.L. Marlowe, A. Puga, Aryl hydrocarbon receptor, cell cycle regulation, toxicity, and tumorigenesis, J Cell Biochem. 96 (2005) 1174–1184. 10.1002/jcb.20656.

[9] A. Hospital, J.R. Goñi, M. Orozco, J.L. Gelpí, Molecular dynamics simulations: advances and applications, Adv Appl Bioinform Chem. 8 (2015) 37–47. 10.2147/AABC.S70333.

[10] E. Yuriev, P.A. Ramsland, Latest developments in molecular docking: 2010-2011 in review, J Mol Recognit. 26 (2013) 215–239. 10.1002/jmr.2266.

[11] J.D. Durrant, J.A. McCammon, Molecular dynamics simulations and drug discovery, BMC Biology. 9 (2011) 71. 10.1186/1741-7007-9-71.

[12] R.P.D. Bank, RCSB PDB - 6TFI: PXR IN COMPLEX WITH THROMBIN INHIBITOR COMPOUND 17, (n.d.). https://www.rcsb.org/structure/6tfi (accessed September 16, 2023).

[13] E. Viesi, D.S. Sardina, U. Perricone, R. Giugno, APDB: a database on air pollutant characterization and similarity prediction, Database. 2023 (2023) baad046. 10.1093/database/baad046.

[14] OPLS4: Improving Force Field Accuracy on Challenging Regimes of Chemical Space | Journal of Chemical Theory and Computation, (n.d.). 10.1021/acs.jctc.1c00302 (accessed September 16, 2023).

[15] R.A. Friesner, R.B. Murphy, M.P. Repasky, L.L. Frye, J.R. Greenwood, T.A. Halgren, P.C. Sanschagrin, D.T. Mainz, Extra Precision Glide: Docking and Scoring Incorporating a Model of Hydrophobic Enclosure for Protein−Ligand Complexes, J. Med. Chem. 49 (2006) 6177–6196. 10.1021/jm051256o.

[16] J.R. Greenwood, D. Calkins, A.P. Sullivan, J.C. Shelley, Towards the comprehensive, rapid, and accurate prediction of the favorable tautomeric states of drug-like molecules in aqueous solution, J Comput Aided Mol Des. 24 (2010) 591–604. 10.1007/s10822-010-9349-1.

[17] K.J. Bowers, D.E. Chow, H. Xu, R.O. Dror, M.P. Eastwood, B.A. Gregersen, J.L. Klepeis, I. Kolossvary, M.A. Moraes, F.D. Sacerdoti, J.K. Salmon, Y. Shan, D.E. Shaw, Scalable Algorithms for Molecular Dynamics Simulations on Commodity Clusters, in: SC ‘06: Proceedings of the 2006 ACM/IEEE Conference on Supercomputing, 2006: pp. 43–43. 10.1109/SC.2006.54.

[18] P. Mark, L. Nilsson, Structure and Dynamics of the TIP3P, SPC, and SPC/E Water Models at 298 K, J. Phys. Chem. A. 105 (2001) 9954–9960. 10.1021/jp003020w.

